# A high-content platform for physiological profiling and unbiased classification of individual neurons

**DOI:** 10.1101/2021.01.25.428085

**Authors:** Daniel M. DuBreuil, Brenda Chiang, Kevin Zhu, Xiaofan Lai, Patrick Flynn, Yechiam Sapir, Brian J. Wainger

## Abstract

High-throughput physiological assays often lose single cell resolution, precluding subtype-specific analyses of neuronal activation mechanism and drug effects. Here, we demonstrate APPOINT, Automated Physiological Phenotyping Of Individual Neuronal Types. This physiological assay platform combines calcium imaging, robotic liquid handling, and automated analysis to generate physiological activation profiles of single neurons at a large scale. Using unbiased techniques, we quantify responses to multiple sequential stimuli, enabling subgroup identification by physiology and probing of distinct mechanisms of neuronal activation within subgroups. Using APPOINT, we quantify primary sensory neuron activation by metabotropic receptor agonists and identify potential contributors to pain signaling. Furthermore, we expand the role of neuroimmune interactions by showing that human serum can directly activate sensory neurons, elucidating a new potential pain mechanism. Finally, we apply APPOINT to develop a high-throughput, all-optical approach for quantification of activation threshold and pharmacologically separate the contributions of distinct ion channel subsets to optical activation.

## Introduction

Modern molecular and genetic tools reveal intricacies of neuronal subtypes in stunning granularity(Hrvatin et al., 2018; Lin et al., 2015; Lubeck and Cai, 2012); however, physiological techniques lag in their ability to provide a combination of large scale and high resolution. Low-throughput, rig-based patch clamp and calcium imaging remain the most common approaches for physiological analyses, due to the wealth of pharmacological and kinetic information generated(Wainger et al., 2015), but cannot capture neuronal diversity or evaluate large compound libraries. Plate reader-based calcium imaging provides high throughput, but only field-wide metrics, and thus can neither distinguish intracellular signal from extracellular noise nor quantify responses among individual cells(Brenneis et al., 2014; Stacey et al., 2018). High-throughput techniques for direct measurement of action potential firing, such as multi-electrode array(Moakley et al., 2019; Wainger et al., 2014a, 2015) or automated patch-clamp recording(Dunlop et al., 2008), lose single cell resolution, require large numbers of cells, or are incompatible with critical primary cell types, such as neurons. A technique that quantifies neuronal activity at large scale while retaining single cell resolution across subtypes would yield substantial benefit for both hypothesis testing and drug screening.

The value of such a tool may be particularly high for identifying new mechanistic targets and treatments for chronic pain, as the lack of adequate treatments underlies the more than 50 million adults in the United States suffering from chronic pain and fuels the ongoing opioid epidemic(Dahlhamer et al., 2018; Kingwell, 2019; Okie, 2010; Yekkirala et al., 2017). Genetic and physiological evidence(Cao et al., 2016; Cox et al., 2006) from rodents and humans supports the therapeutic potential of blocking activation of nociceptors, first-order sensory neurons that initiate acute and chronic pathological pain(Basbaum et al., 2009; Hucho and Levine, 2007). Nociception can be interrogated *in vitro* using primary rodent, human post-mortem(Valtcheva et al., 2016), and human stem cell-based models(Schwartzentruber et al., 2018; Wainger et al., 2015), but two considerations demonstrate the need for physiological techniques with improved throughput and single cell resolution. First, independent subgroups of sensory neurons mediate and distinguish diverse sensory percepts, including pain, pleasant touch, and itch(Albisetti et al., 2019; Huang et al., 2018; Li et al., 2016; Seal et al., 2009; Usoskin et al., 2014). Indeed, independent transcriptomic analyses have revealed more than 10 distinct types of mouse sensory neurons(Li et al., 2016; Usoskin et al., 2014). Second, individual receptors and ion channels transduce signals in a context-dependent manner based on the cellular and molecular environment of distinct neuronal subtypes(Hill et al., 2018; Rush et al., 2006). These considerations build a strong case for a phenotypic approach to target and drug identification using sensory neurons with sufficient scale to assess effects in both common and rare nociceptor types.

Here, we use a high-throughput platform for Automated Physiological Phenotyping Of Individual Neuronal Types (APPOINT, Supplementary Figure 1) to quantify the activation of primary and human induced pluripotent stem cell (iPSC)-derived neurons in response to automated application of chemical and optogenetic stimuli. APPOINT utilizes unbiased hierarchical clustering and random forest machine learning approaches to distinguish positive and negative responses to each of several sequential stimuli, and then uses the response patterns for each individual cell to assemble its physiological phenotype. We apply APPOINT to quantify the extent and breadth of activation of sensory neuron subtypes by a panel of ionotropic and metabotropic receptor agonists, demonstrate a new potential pain mechanism by quantifying sensory neuron activation by human blood serum, and develop an approach for measuring optogenetic activation threshold with pharmacological validation of contributions from physiologically-relevant ion channels. Thus, APPOINT is a flexible platform that enables diverse high-throughput physiological assays for neuronal excitation and holds promise for identification and testing of novel neuronal activation mechanisms as well as development of non-opioid therapies for pain.

## Results

### Setup of high-throughput calcium imaging assay

We first validated primary sensory neuron isolation and plating protocols to ensure reproducibility within and among neuronal batches. We isolated primary mouse dorsal root ganglia and plated dissociated neurons into 96-well plates coated with poly-D-lysine and laminin in the center of each well (Figure 1a). Cells were imaged in a single central field for each well (Figure 1b) and neuronal cell bodies were counted automatically using a custom analysis script (see Materials and Methods), removing any potential bias from site or cell selection. Across a sample of 859 wells in 13 independent batches, each corresponding to an independent imaging plate containing neurons from one or two mice, we analyzed an average of 137.2 ± 3.5 cells/well, or 13,000 cells/96 well batch (Figure 1c-1d). The mean number of cells/well by batch ranged from 61 to 264 cells/well. Within batches, the coefficient of variation was low and few wells had fewer than 20 cells (Supplementary Figure 2). We did not observe a significant difference between edge and center wells in neuronal density (main effect of edge by univariate ANOVA, n=316 edge wells/543 center wells, F=0.49, p=0.484); therefore, we continued to use all wells per plate. Traditional rig-based experiments generally use fewer than 100 cells/group(Bataille et al., 2020; Baykara et al., 2019; Tonello et al., 2017); thus, using the lowest average cell count per well, we would only require two wells per group to surpass current typical sample sizes.

**Figure 1.**
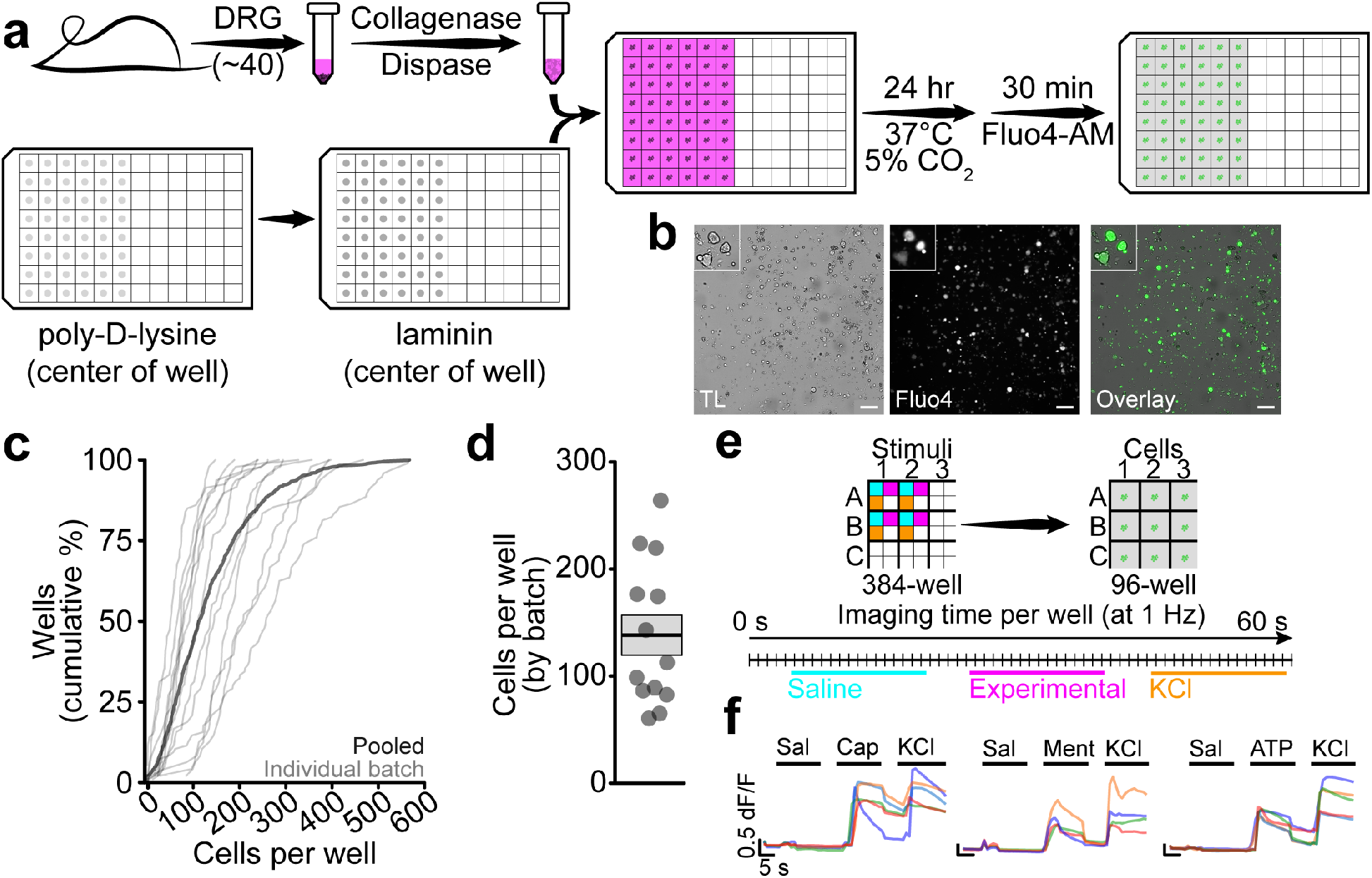
Sensory neuron isolation, plating, and automated stimulation. **a**, Overview of DRG isolation and plating in 96-well format. **b**, Example images of dissociated DRG neurons visualized by transmitted light (TL, grey, left) and Fluo4 (green, middle) along with overlay (right). Scale bar indicates 200 μm and inset shows cellular detail. **c**, Cumulative distribution of cells per well for each of 13 plates individually (thin lines) and pooled (thick line). **d**, Average number of cells per well within each plate (points), along with mean and SEM (box). **e**, Schematic overview of liquid handling and imaging protocol for sequential stimuli. **f**, Example traces from individual cells showing consecutive activation by one of capsaicin (200 nM, left), menthol (300 μM, middle), or meATP (250 μM, right) and KCl (100 mM) (scale bars 5 s and 0.5 dF/F).

Application of consecutive chemical stimuli to individual cells in rig-based microperfusion systems has allowed for cell type-specific analysis of neuronal function(Bautista et al., 2006; Caterina et al., 2000; Chen et al., 1995; Hill et al., 2018; Yin et al., 2018). Therefore, we developed a stimulation protocol to apply multiple stimuli to cells in each imaging well, with the goal of facilitating physiological phenotyping of functional cell types. Three chemical stimuli were applied at 20 s intervals from 384-well plates to each well of cells in 96-well plates (Figure 1e). This configuration allows up to 8 chemical stimuli using two stimulus plates. Each cell was exposed to negative and positive control stimuli—saline and high K^+^—as well as capsaicin, menthol, or αβ-methyleneadenosine 5’-triphosphate (meATP), which activate TrpV1, TrpM8, and P2×3 receptors, respectively (Figure 1f). Within individual cells, we observed large amplitude responses to both K^+^ and all three ionotropic receptor agonists.

### Unbiased quantification of high-throughput calcium imaging using machine learning techniques

Current methods for collecting and analyzing calcium imaging data using rig-based platforms require substantial experimenter input for selecting sites for imaging, cells for analysis, and amplitude thresholds for distinguishing responsive and non-responsive cells(Barabas et al., 2012; Than et al., 2013). Each of these inputs increases the likelihood of bias, reduces reproducibility across groups, and, at the same time, decreases throughput. Using our approach, we have already removed empirical site and cell selection via our automated imaging protocol and we next developed an unbiased pipeline for quantification of neuronal activation.

We used a large dataset of 58,635 neurons from 10 batches of mouse primary sensory neurons, with each neuron stimulated by three stimuli, including saline, one nociceptor-specific ionotropic receptor ligand (capsaicin, menthol, or meATP), and high K^+^. Across all neurons, we isolated the responses to each stimulus, so that each could be classified as positive or negative independently, and measured the peak response amplitude and maximum rate of rise of Fluo4 intensity. We initially pooled together all responses to saline or high K+, ignoring responses to capsaicin, menthol, and meATP, within each batch and used agglomerative hierarchical clustering to form three clusters of responses. We examined traces sorted into each cluster and found that, in every batch tested, a single cluster contained primarily negative responses and both other clusters contained almost exclusively positive responses (Figure 2a). All responses sorted into the cluster exhibiting the smallest average amplitude and shallowest average slope were considered negative, and responses sorted into the other clusters were considered positive. When we examined cumulative distributions of the peak response amplitudes for responses classified as positive or negative (Figure 2b), we observed overlap between the two distributions; however manual qualitative spot checks of responses classified as positive and negative within the low amplitude range (Figure 2c) revealed striking differences in the dynamics of the calcium response, with the negative responses visually indicative of stimulus artifacts resulting from liquid injection. Notably, use of a fixed threshold, as is currently the standard approach to such analyses(Blanchard et al., 2015; Perner et al., 2020), which would have classified these artifacts as positive responses, and thus would have resulted in increased false positive responses.

**Figure 2.**
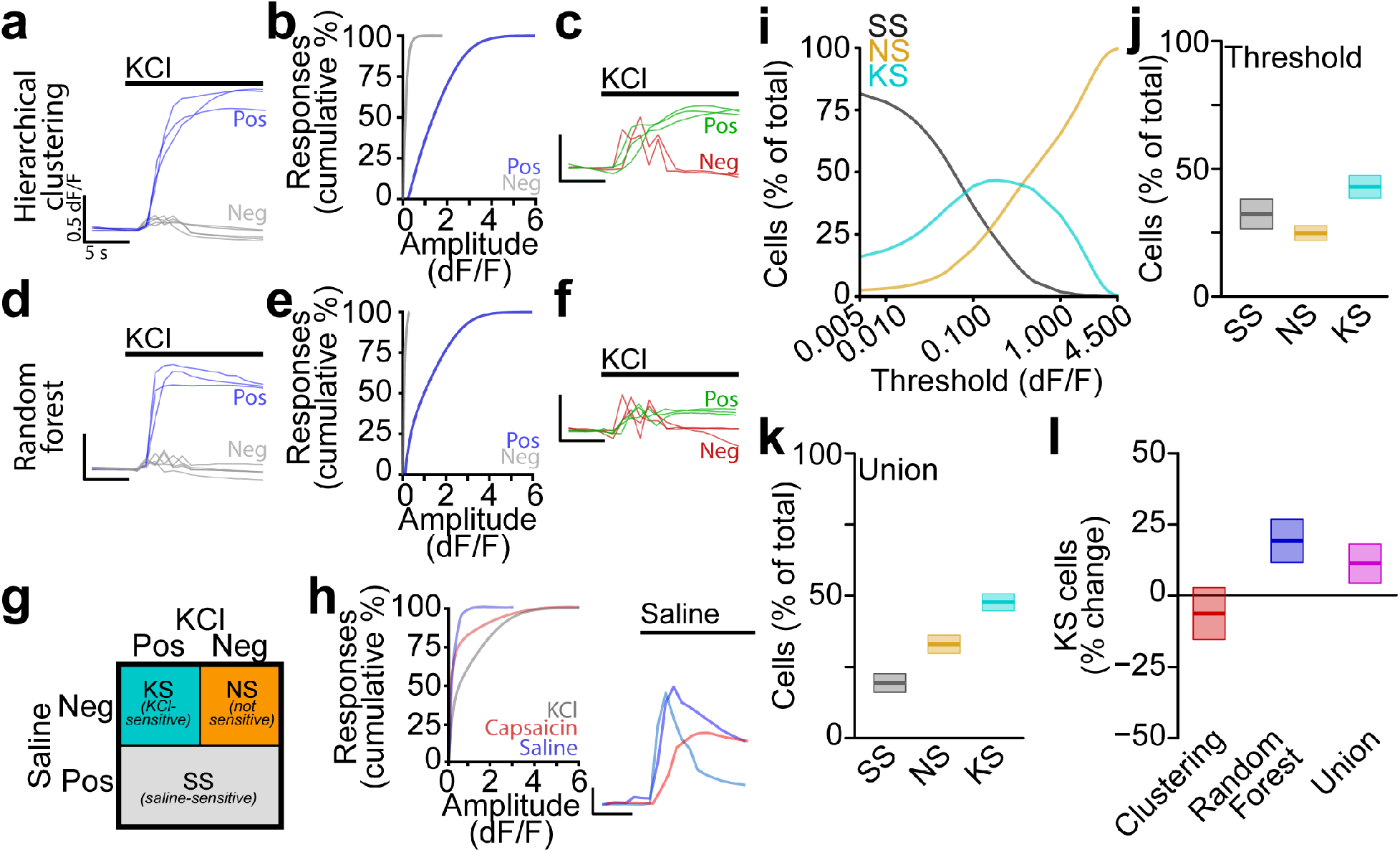
Machine learning-enabled classification of automated calcium imaging responses. **a**, Example traces of KCl-dependent calcium responses classified as positive (blue) and negative (grey) by hierarchical clustering (scale bars 5 s and 0.5 dF/F). **b**, Cumulative distributions of positive (blue) and negative (grey) response amplitudes. **c**, Example traces of small amplitude KCl-dependent calcium responses classified as negative (red) and positive (green) by hierarchical clustering (scale bars 5 s and 0.5 dF/F). **d**, Example traces of KCl-dependent calcium responses classified as positive (blue) and negative (grey) by random forest model (scale bars 5 s and 0.5 dF/F). **e**, Cumulative distributions of positive (blue) and negative (grey) response amplitudes. **f**, Example traces of small amplitude KCl-dependent calcium responses classified as negative (red) and positive (green) by random forest model (scale bars 5 s and 0.5 dF/F). **g**, Cell classification scheme based on coordinated responses to saline and KCl stimuli. **h**, Cumulative distributions of response amplitudes (including both positive and negative, left) for saline (blue), capsaicin (red), and KCl (grey) along with example traces (right, colors indicate individual cells, scale bars 5 s and 0.5 dF/F) **i**, Percentage of total cells classified as saline-sensitive (SS, grey), not sensitive (NS, orange), and KCl-sensitive (KS, cyan). **j**, Percentage of cells classified as SS (grey), NS (orange), and KS (cyan) using standard threshold of 0.15 dF/F. **k**, Percentage of cells classified as SS (grey), NS(orange), and KS(cyan) using the union of hierarchical clustering and random forest classifiers. **l**, Percent change from threshold-based analysis in percentage of KS cells using hierarchical clustering (red) or random forest (blue) classifiers alone or the union of both approaches (magenta).

We then repeated this analysis including responses to capsaicin, menthol, and meATP along with saline and high K^+^ (Supplementary Figure 3). Positive responses to each of the three ionotropic receptor activators qualitatively matched responses to high K^+^ and were easily distinguished from negative responses. Furthermore, as noted above, we observed a similar degree of overlap in the amplitude of positive and negative responses, with large negative responses resembling smaller positive responses. Thus, hierarchical clustering permits unbiased classification of calcium responses to several independent chemical stimuli and decreases false-positive calls by use of additional response properties beyond peak amplitude.

Hierarchical clustering is an unsupervised machine learning approach, and we thought that a supervised approach might provide a robust complement to validate clustering-based classification. Therefore, we trained random forest models, one for each neuronal batch, to identify positive and negative responses. In addition to measurements of peak amplitude and maximum rate of change of Fluo4 intensity, we included raw baseline and peak fluorescence intensities as additional features. In total, we trained each model with less than 5% of responses from each batch and then predicted the classification for the remaining 95% of responses. After classification, we examined responses classified as positive and negative and verified the classification qualitatively for high K^+^ and ionotropic receptor responses (Figure 2d, Supplementary Figure 3). There was much less overlap between cumulative amplitude distributions for positive and negative responses (Figure 2e) compared to the hierarchical clustering method, and large amplitude stimulus artifacts were again properly classified as negative (Figure 2f). To assess the robustness of our response models, we also tested whether classifiers trained on individual batches could correctly identify responses when tested on each other batch. For every pair of batches, considering the within-batch classification as ground truth for each response, cross-batch model accuracy was significantly above chance (87.6±0.6%, Supplementary Table 1), indicating robust response classification by the models across independent batches.

These approaches use multiple features of each calcium response and rely on distinct methodologies for classifying responses; however, they were remarkably consistent. Across all 117,270 individual stimulus responses, the two approaches agreed on 88.4% of responses, with a sensitivity of 82.3% and specificity of 96.6%. For the remainder of the study, we considered all responses classified as positive by both approaches as true positive responses.

### Application of unbiased response classifications for phenotyping individual cells

Classification of individual responses as positive or negative is critical for accurate quantification of cell types, but the overarching goal of this platform is to physiologically classify cells based on coordinated responses to multiply stimuli. Thus, to address this, we sorted a sample of 58,635 neurons from 10 batches of mouse primary sensory neurons into one of three groups based on their responses to positive and negative stimuli, namely high K^+^ and saline, respectively (Figure 2g). All saline responsive cells, regardless of K^+^ sensitivity, were grouped together (SS, saline sensitive), whereas saline-insensitive cells were classified as either not responsive (NS) or K^+^ sensitive (KS). Positive responses to saline were relatively rare compared with capsaicin and high K^+^, but large amplitude responses were clearly distinguishable from stimulus artifacts or noise (Figure 2h). Mechanosensitive channels likely contribute to these saline-dependent responses, but the multitude of potential mechanisms complicates investigation of their physiological relevance(Beaulieu-Laroche et al., 2020; Coste et al., 2012; Dong et al., 2015; Kanda et al., 2019; Murthy et al., 2018). To compare our method with traditional threshold-based approaches, we classified cells using a wide range of threshold values and quantified the relative proportion of cells in each category (Figure 2i). At low threshold values, most cells were classified as SS, whereas, at higher thresholds, most cells were NS. Between these threshold extremes, KS cells comprised the largest group.

Using a typical amplitude threshold of 0.15 dF/F, we observed that roughly equal proportions of cells were sorted into each category (SS 32.3±5.8% per batch; NS 24.8±2.9% per batch; KS 42.9±4.5% per batch; Figure 2j). The hierarchical clustering approach generated a similar pattern (SS 17.1±3.1% per batch; NS 43.0±3.3% per batch; KS 39.8±3.2% per batch; Supplementary Figure 4a), but, using the random forest approach or the union of the two approaches, we observed a clear increase in cells classified as KS relative to SS or NS (Random Forest: SS 21.3±3.3% per batch, NS 27.6±2.4% per batch, KS 51.1±3.3% per batch; Figure Supplementary Figure 4b; Union: SS 23.2±3.7% per batch, NS 26.8±2.4% per batch, KS 50.0±3.5% per batch; Figure 2k). Furthermore, when directly comparing the three unbiased approaches to the conventional threshold approach by batch, we observed that the random forest and union approaches categorized more cells as KS (Random forest: 19.0±7.7% increase; Union: 16.5±8.1% increase, n=10 batches, Figure 2l) and fewer cells as SS (Random forest: 34.0±10.1% decrease; Union: 28.0±11.3% decrease, n=10 batches, Supplementary Figure 4c), with no change in NS cells (Random forest: 11.4±9.6% increase; Union: 8.0±9.8% increase, n=10 batches, Supplementary Figure 4d). These results demonstrate that unbiased hierarchical clustering and random forest machine learning classifiers can robustly distinguish positive and negative calcium responses in individual neurons and enable high-throughput classification of neurons based on physiological responses to consecutive stimuli.

### Neuronal subtype determination by physiological activation of cell type-specific ionotropic receptors

We next applied to APPOINT to quantify neuronal activation in response to increasing doses of K^+^ or ionotropic receptor agonists. Increasing concentrations of K^+^ elicited progressively larger-amplitude calcium responses (Figure 3a) and recruited larger populations of neurons (Figure 3b). The percentage of neurons that was activated by K^+^ reached a plateau of 57.8±5.5% around 35 mM K^+^, with an EC50 of 16.4±3.6 mM K^+^. We also applied capsaicin, menthol, and meATP at five doses each and observed robust activation (Figure 3c-d). Both response amplitude distributions and neuron recruitment showed strong dose sensitivity, with increasing doses yielding larger responses and increased numbers of responding neurons (number of responsive cells assessed by univariate ANOVA, n=6 batches, capsaicin: F=36.76, p=1.3*10^−7^; menthol: F=60.06, p=2.2*10^−10^; meATP: F=6.129, p=0.016).

**Figure 3.**
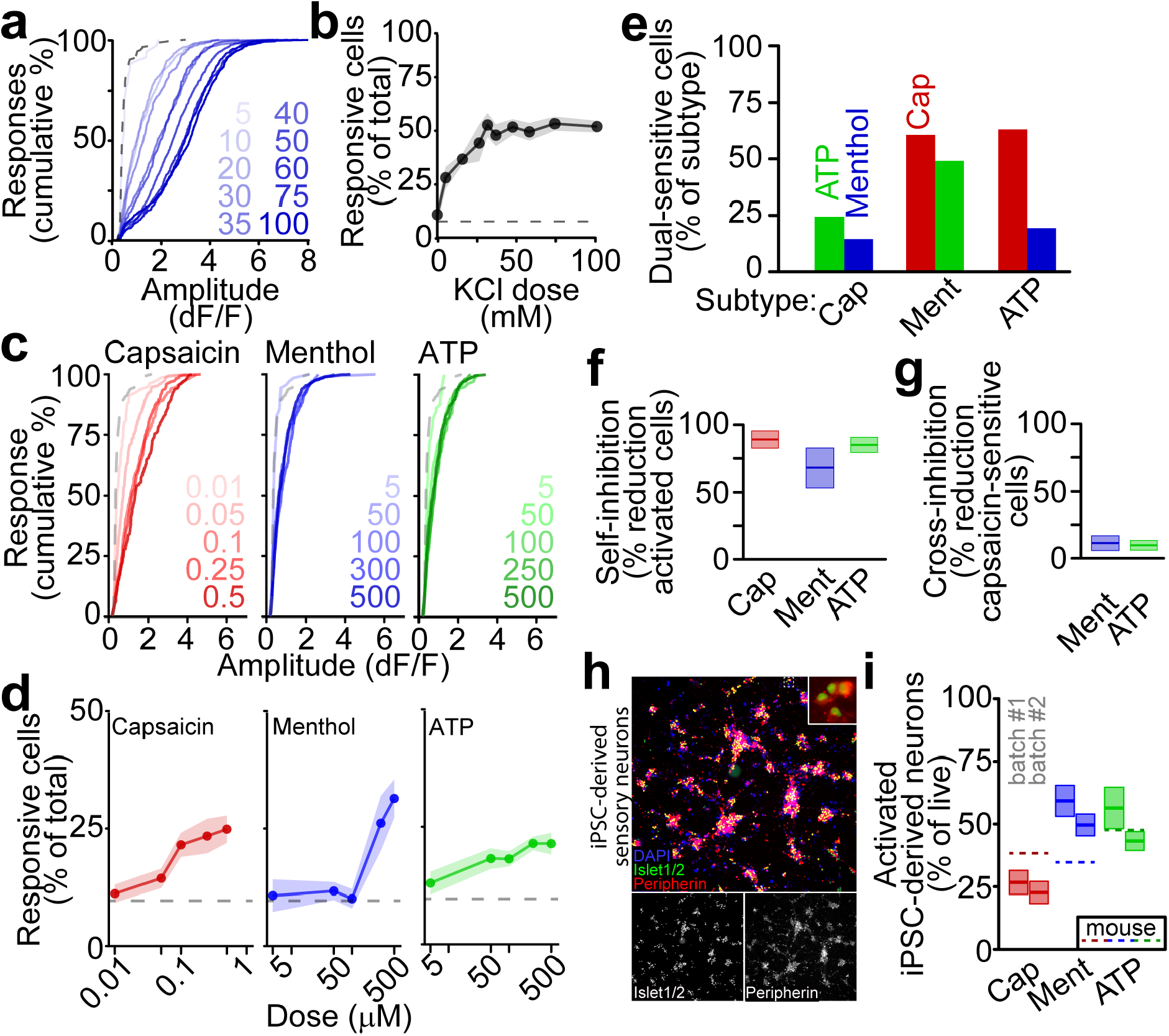
Quantification of sensory neuron subtypes by ionotropic receptor activation. **a**, Cumulative distributions of peak amplitudes following stimulation by increasing doses of KCl (mM, blue) along with saline (dashed grey). **b**, Percentage of neurons activated by increasing doses of KCl, as mean (point/line) and SEM (shaded area). **c**, Cumulative distributions of positive response amplitudes following stimulation by increasing doses (in μM) of capsaicin (red), menthol (blue), and meATP (green), along with saline (dashed grey line). **d**, Percentage of cells activated by increasing doses of capsaicin (red), menthol (blue), and meATP (green), along with saline (dashed grey line). **e**, Percentage of nociceptor subtypes functionally co-expressing other subtype-specific receptors. **f**, Self-inhibition of capsaicin (red, 100 nM), menthol (blue, 300 μM), or meATP (green, 250 μM) activation by prior stimulus as mean and SEM. **g**, Cross-inhibition of capsaicin (100 nM) responses by prior stimulation with menthol (blue, 300 μM) or meATP (green, 250 μM) as mean and SEM. **h**, iPSC-derived sensory neurons at 5 weeks in culture stained with DAPI (blue), anti-Islet1/2 (green), and anti-Peripherin (red). Overlay inset shows cellular detail. **i**, Percentage of live iPSC-derived neurons activated by ionotropic receptor agonists at 9 weeks in culture as mean and SEM for two independent differentiations. Dashed lines show average activation of primary mouse sensory neurons.

Although these agonists identify broad subpopulations of sensory neuron types, there is disagreement regarding the extent of their functional co-expression(Li et al., 2016; Usoskin et al., 2014). Therefore, we quantified functional co-expression of all three pairwise combinations of receptors. We applied each pair of agonists, with equal wells for each pair and each unique stimulus order, then quantified the number of neurons responding to both stimuli (Figure 3e). Interestingly, we observed co-expression of each receptor pair. Capsaicin-sensitive cells were rarely activated by meATP (24.4%) or menthol (14.5%), whereas menthol activated-cells were frequently activated by capsaicin (60.7%) or meATP (49.1%). Although we observed positive responses to each stimulus, whether applied first or second, we also quantified inhibition resulting from consecutive application of stimuli (Figure 3f-g). As expected, activation by all three stimuli was strongly inhibited by prior application of the same stimulus (capsaicin: 89.0±6.4%; menthol: 68.0±14.7%; meATP: 85.0±5.7%; Figure 3f); however, activation by capsaicin was very weakly inhibited by prior application of either menthol (10.1.0±6.4%) or meATP (9.2±4.1%). Thus, although there is the potential for cross-stimulus inhibition, as occurs with conventional rig-based calcium imaging platforms as well, APPOINT permits robust unbiased quantification of interference.

Although mouse primary sensory neurons provide a useful and easy source of neurons for analysis, higher-throughput applications of APPOINT will require expandable cell-types, such as human induced pluripotent stem cell (iPSC)-derived neurons. We next quantified subtypes of iPSC-derived sensory neurons using physiological responses to cell type-specific activators. Sensory neurons were differentiated from a human iPSC line according to an established protocol(Schwartzentruber et al., 2018) and expression of the sensory neuron markers Islet and Peripherin was verified at 5 weeks in culture (Figure 3h). Across two independent differentiation batches of iPSC-derived neurons, positive responses to each ionotropic receptor agonist were observed after 9 weeks in culture, with similar proportions of each nociceptor subtype observed across batches (Figure 3i, for batch #1/batch #2, n=8 wells/12 wells per stimulus, capsaicin: 26.7±4.8%/22.7±4.4% of live cells, p=0.9996; menthol: 59.3±6.2%/49.6±4.4% of live cells, p=0.927; meATP: 56.4±8.3%/43.2±3.8% of live cells, p=0.714; significance assessed as batch*stimulus interaction by univariate ANOVA followed by Tukey HSD correction for multiple comparisons). Comparing receptor expression in iPSC-derived and primary mouse neuronal cultures, iPSC-derived sensory neuron cultures generated notably fewer capsaicin-sensitive and more menthol-sensitive cells, a common trend observed in cultures of iPSC-derived sensory neurons(Blanchard et al., 2015; Chambers et al., 2012; Nickolls et al., 2020).

### Systematic identification of metabotropic receptors mediating calcium flux and pain

Metabotropic receptors contribute to inflammatory and neuropathic pain by modulating sensory neuron excitability through activation of intracellular signaling cascades and modulation of gene expression(Eskander et al., 2015; Gibbs et al., 2007; Hendrich et al., 2012; Melemedjian et al., 2010; Mendieta et al., 2016). Most studies focus on sensitization of nociceptors by inflammatory mediators, but some mediators, such as bradykinin and prostaglandin E2 (PGE2), directly activate calcium flux and cause pain in humans and rodents(Hong and Abbott, 1994; Linhart et al., 2003; Liu et al., 2010; Mørk et al., 2003; Oh et al., 2001). The broad ability of inflammatory mediators to activate sensory neurons has not been addressed systematically; thus, we applied APPOINT to quantify nociceptor activation by a pool of metabotropic receptor agonists.

We performed unbiased analysis of publicly available transcriptomic datasets to identify metabotropic receptors potentially involved in pain signaling based on high expression levels in DRG (GSE63576(Li et al., 2016)), enriched expression in DRG (GSE10246(Lattin et al., 2008)) and nociceptors (GSE55114(Chiu et al., 2014)), and changes in expression levels in the spared nerve injury model of neuropathic pain (GSE89224(Cobos et al., 2018)). We selected all receptors with commercially available and well-validated small molecule activators and obtained a pool of 18 receptor-agonist pairs (Supplementary Table 2). We then tested three doses for each compound (listed in Supplementary Table 2) and observed a wide range of activation profiles (Figure 4a). We included the inflammatory mediators bradykinin and prostaglandin E2 (PGE2) as positive control stimuli, indicating levels of activation produced by compounds known to elicit pain(Hong and Abbott, 1994; Linhart et al., 2003; Mørk et al., 2003). Bradykinin was the most effective agonist (Figure 4b), but several other agonists, including neuropeptide Y, sulprostone, and L-054,264, activated at least as many sensory neurons as PGE2. Every agonist tested activated more neurons than saline; however, several candidates with prominent roles in pain, including nerve growth factor(Denk et al., 2017) (NGF) and interleukin 6(Zhou et al., 2016), directly activated surprisingly small subpopulations. Calcium flux in these neurons is likely due to activation of intracellular signaling leading to intracellular calcium release, which may be related to neuronal sensitization(Alkhani et al., 2014; Cesare et al., 1999; Coderre, 1992).

**Figure 4.**
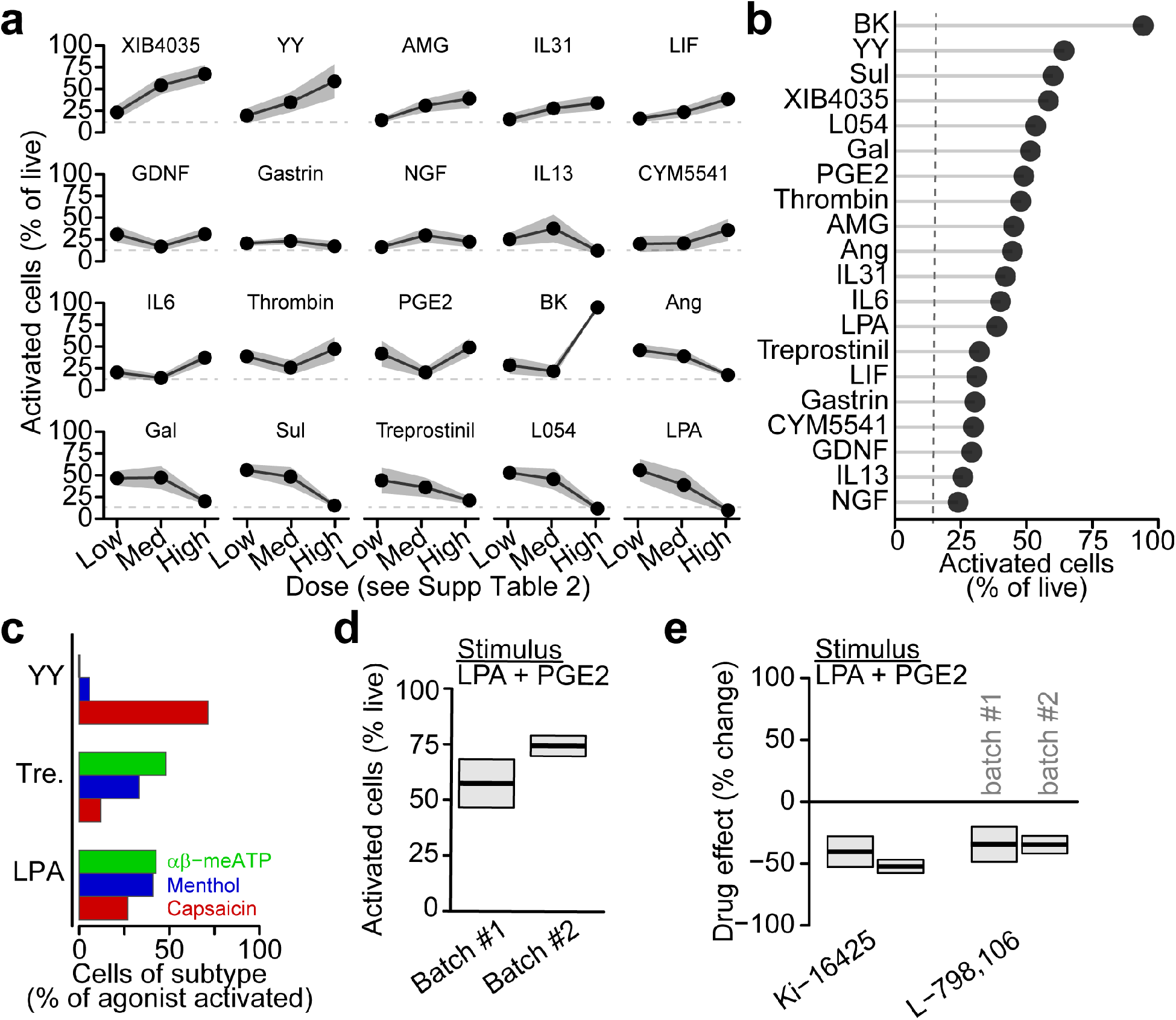
Metabotropic receptor activation elicits calcium flux in primary mouse sensory neurons. **a**, Percentage of live mouse primary sensory neurons activated by metabotropic receptor agonists (Supplementary Table 2) as mean (points/lines) and SEM (shaded area). Dashed grey line indicates activation by saline. **b**, Rank ordering of metabotropic receptor agonists at single dose activating highest percentage of neurons. **c**, Activation of nociceptor subtypes by selected metabotropic receptor agonists. **d**, Percentage of live primary sensory neurons activated by a cocktail of metabotropic receptor agonists, LPA (200 μM) and PGE2 (1 mM) as mean and SEM of two independent batches. **e**, Percent change in cocktail-activated cells by receptor-specific antagonists, Ki-16425 (LPA3, 10 μM), L-798,106 (EP3, 10 μM), relative to vehicle as mean and SEM.

We then examined the specific types of sensory neurons that were activated by a subset of these metabotropic receptor agonists: neuropeptide Y, treprostinil, and lysophosphatidic acid (LPA). Neuropeptide Y selectively activated capsaicin-sensitive nociceptors, whereas both treprostinil and LPA activated all three subtypes of sensory neurons (Figure 4c).

To assess blockers of metabotropic receptor activation, we first combined LPA and PGE2 into a single stimulus cocktail and quantified inhibition by blockers of each target receptor, LPA-3 and EP3, independently. Across two independent batches of primary mouse sensory neurons, we observed consistent neuronal activation by the LPA+PGE2 cocktail (Figure 4d, for batch #1/batch #2, n=16 wells/14 wells, mean and SEM: 57.5±10.8%/74.5±4.6% of live cells, main effect of batch assessed by univariate ANOVA, F=1.876, p=0.182) and pretreatment with either Ki-16425, an inhibitor of LPA-1 and LPA-3 receptors(Velasco et al., 2017), or L-798,106, an inhibitor of EP3 receptors(Shridas et al., 2014), decreased activation relative to vehicle (Figure 4e, for batch #1/batch #2, n=16 wells/batch/drug, Ki-16425: 40.7±12.6% decrease, one-sample t-test, t=-3.244, p=0.005/52.7±5.5% decrease, one-sample t-test, t=-9.57, p=8.9*10^−8^; L-798,106: 34.6±14.5% decrease, one-sample t-test, t=-2.389, p=0.030/35.0±7.1% decrease, one-sample t-test, t=-4.906, p=0.0002). Furthermore, within each drug, we did not observe a significant difference in the magnitude of inhibition across batches (Ki-16425: n=16 wells/batch, p=0.853; L-798,106: n=16 wells/batch, p=0.999; significance assessed as drug*batch interaction by univariate ANOVA with Tukey HSD post-hoc correction for multiple comparisons). We conclude that many metabotropic receptors directly elicit calcium flux in subgroups of sensory neurons, demonstrating the power of this approach to identify novel activators and inhibitors of metabotropic receptor-mediated calcium flux in cellular context.

### Human serum directly activates calcium flux in primary sensory neurons

Increasing evidence suggests a dynamic interaction between vascular permeability and pain via inflammatory mediators released locally by invading immune cells(Chiu et al., 2012; Pinho-Ribeiro et al., 2017); however, we hypothesized that blood itself may directly elicit calcium flux in sensory neurons. To test this, we obtained serum samples from 9 healthy volunteer participants (Figure 5a) and used APPOINT to quantify the effects of serum application on primary mouse sensory neurons. Serum elicited large amplitude calcium responses that qualitatively resembled responses to ionotropic and metabotropic receptor agonists (Supplementary Figure 5a). Initially, we quantified responses in all neurons, regardless of sensitivity to saline and observed strong dose-response relationships for both response amplitude (Supplementary Figure 5b) and the percent of cells activated by serum (Supplementary figure 5c). Due to the substantial activation even at 1000-fold dilution, we then discarded from analysis all cells that responded saline, which may respond non-specifically to liquid handling. Nevertheless, we again observed robust dose-dependence for both response amplitudes (Figure 5b) and percent of cells activated (Figure 5c). Serum elicited responses of similar amplitude (Figure 5d) in sensory neurons sensitive to capsaicin, menthol, and meATP and the percentage of serum sensitive cells was similar across all three subtypes (Figure 5e, capsaicin: 20.6±2.2%, n=31 wells across 3 batches; menthol: 37.9±4.7%, n=18 wells across 3 batches; meATP: 24.8±4.7%, n=18 wells across 2 batches). We then sought to identify the active components of serum that contribute to neuronal activation by 1) heat inactivation of the complement system(Soltis et al., 1979), 2) digestion of proteins using proteinase K, and 3) removal of the 12 most abundant serum proteins (see Materials and Methods) by spin column filtration (Figure 5f-g). Heat inactivation reduced neither the amplitude (Figure 5f) nor the percentage of cells activated (Figure 5g, original: 23.4±1.3%, n=78 wells across 3 batches; heat inactivated: 25.8±2.2%, n=26 wells across 1 batch), indicating no contribution of complement components. By contrast, both proteinase K treatment and removal of the top 12 abundant serum proteins decreased the amplitude of serum-mediated calcium responses and the percentage of activated neurons (proteinase K: 14.3+1.2%, n=52 wells across 2 batches; filtered: 8.3±0.9%, n=82 wells across 2 batches). Finally, we quantified activation by serum samples from each volunteer individually (Figure 5h). Serum-mediated activation was observed using all nine samples, indicating that the ability to activate sensory neurons is a general feature of human blood serum and may suggest a potential new pain mechanism.

**Figure 5.**
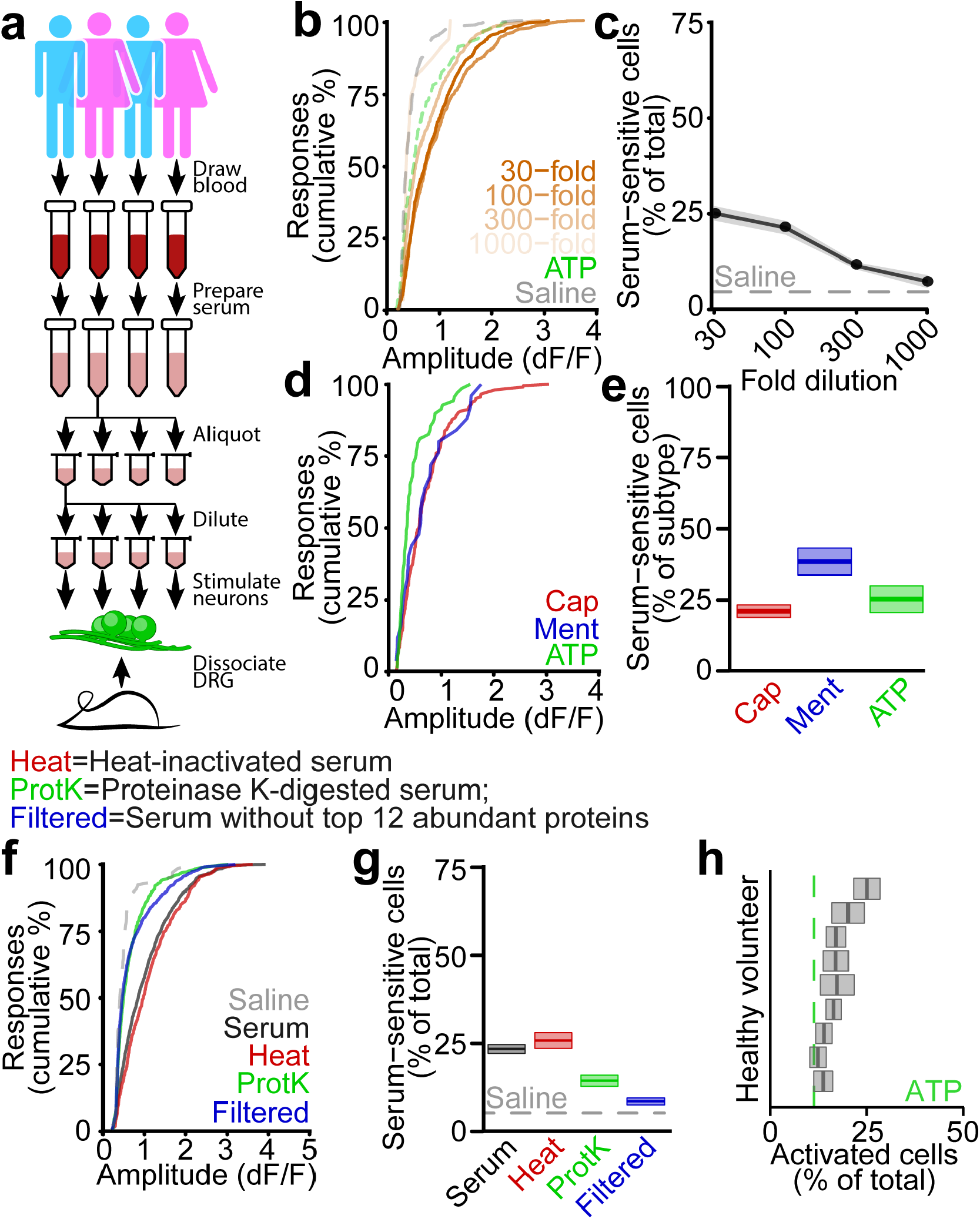
Activation of primary mouse nociceptors by human blood serum. **a**, Protocol for stimulating primary mouse sensory neurons with human serum. **b**, Cumulative distributions of positive response amplitudes following stimulation by serum dilutions (as fold dilution, orange), along with meATP (dashed green line) and saline (dashed grey line). Analysis excludes cells sensitive to initial saline stimulus. **c**, Dose-response curve of sensory neuron activation by serum as mean (points/lines) and SEM (shaded area), along with saline-dependent activation (dashed line). Analysis excludes cells sensitive to initial saline stimulus. **d**, Cumulative distributions of amplitudes of serum-dependent calcium responses in capsaicin-sensitive (red), menthol-sensitive (blue), or meATP-sensitive (green) sensory neurons. **e**, Percentage of capsaicin-sensitive (red), menthol-sensitive (blue), and meATP-sensitive (green) sensory neurons activated by serum (300 fold dilution) as mean and SEM. **f**, Cumulative distributions of amplitudes of serum-dependent calcium responses following heat inactivation (Heat, red), proteinase K digestion (ProtK, green), or spin column removal of top 12 abundant proteins (Filtered, blue), relative to unmodified serum (black) and saline (grey). **g**, Percentage of sensory neurons activated by unmodified serum (black), heat inactivated serum (Heat, red), proteinase K digested serum (ProtK, green), or serum following abundant protein removal (Filtered, blue), along with saline activation, as mean and SEM. **h**, Percentage of live primary mouse sensory neurons activated by serum samples from 9 healthy volunteers (300 fold dilution) as mean and SEM, along with meATP (250 μM, dashed green line).

### A high-throughput platform for quantification and pharmacological modulation of activation threshold

Activation of ionotropic or metabotropic receptors tests sensitivity to specific ligands. By contrast, basal excitability is ligand-independent, reflecting the cohort of functional ion channels, their modulation states, and propensity to conduct current. Increased neuronal excitability is a pathognomonic feature of pathological pain states(Hucho and Levine, 2007; Ma et al., 2006) as well as other neurological diseases(Devinsky et al., 2013; Geevasinga et al., 2016; Ratté and Prescott, 2016). Therefore, we combined optogenetics and APPOINT to develop an all-optical approach to quantify activation threshold. We generated transgenic mice expressing ChR2-EYFP in Trpv1-lineage neurons by crossing Cre-dependent ChR2-EYFP reporter mice(Madisen et al., 2012) with a Trpv1-Cre mouse strain(Cavanaugh et al., 2011a, 2011b)(Figure 6a) and monitored intracellular calcium using a red-fluorescent calcium-sensitive dye, CalBryte-630AM(Yang et al., 2019).To activate neurons, we applied five stimulus trains of 1 ms light pulses, increasing the number of pulses with each successive stimulus train (Figure 6b). We observed positive responses to at least one of the five stimulus trains in 6.4±0.5% of all neurons (n=28 wells across 6 independent batches) with an average amplitude of 0.301±0.003 dF/F (n=1,671 responses). The percentage of light-sensitive cells increased with the number of stimuli per train (Figure 6c), as did the amplitude of positive responses (Figure 6d). To confirm the dependence of these responses on ChR2-EYFP expression, we applied three different antisense oligonucleotides (ASOs, 10 μM) targeting the ChR2-EYFP gene, along with a scrambled control ASO, to dissociated sensory neurons. Treatment for seven days reduced the percentage of light-sensitive cells relative to control ASO-treated neurons for two of the ChR2-EYFP targeting ASOs (ChR1: 81.6±6.7% decrease, n=10 wells across 2 batches; ChR2: 53.1±7.1% decrease, n=10 wells across 2 batches), with no effect of the third (ChR3: 4.0±13.2% decrease, n=10 wells across 2 batches; Figure 6e-f). None of the ASO treatments affected the amplitude of light-activated calcium responses (Supplementary Figure 6), indicating that, when the ASOs were taken into cells, there was sufficient down-regulation of ChR2-EYFP mRNA and protein turnover to effect a profound decrease of light-dependent activation.

**Figure 6.**
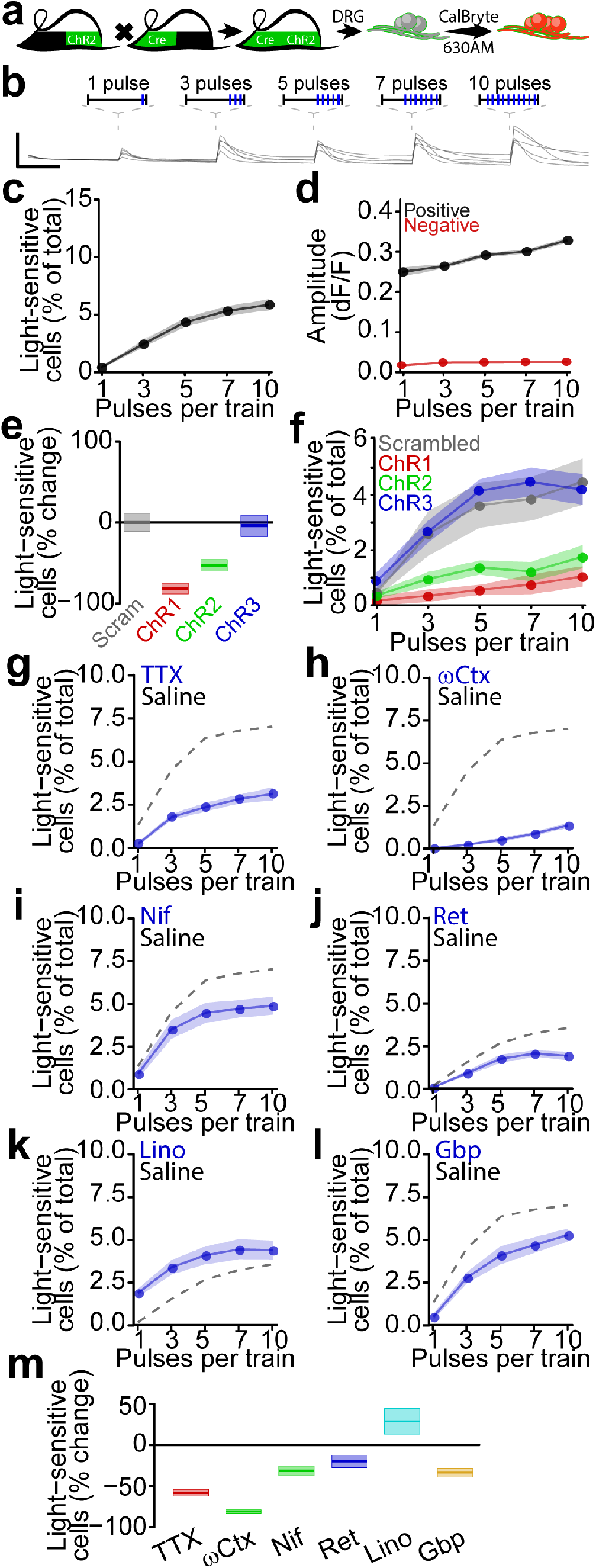
Development of high-throughput activation threshold analysis. **a**, Scheme for calcium imaging of ChR2-expressing primary mouse sensory neurons. **b**, Example traces of calcium flux in individual primary mouse sensory neurons (lines) resulting from optogenetic activation (scale bars 0.5 dF/F and 5 s). **c**, Percentage of cells responding to each optical stimulus train as mean (lines/points) with SEM (shaded). **d**, Average response amplitude for positive (black) and negative (red) responses to each optical stimulus train as mean (lines/points) with SEM (shaded). **e**, Percent change in light-sensitive sensory neurons following seven-day treatment by ChR2-targeting ASOs (red, green, blue, 10 μM) or scrambled control ASO (grey, 10 μM) as mean and SEM. **f**, Percentage of cells responding to each optical stimulus train following seven-day treatment by ChR2-targeting ASOs (red, green, blue) or scrambled control ASO (grey) as mean (point/line) and SEM (shaded). **g-l**, Inhibition of optogenetic activation by tetrodotoxin (**g**, TTX, 300 nM), ω-conotoxin MVIIA (**h**, ωCtx, 3 μM), nifedipine (**i**, Nif, 10 μM), retigabine (**j**, Ret, 10 μM), linopirdine (**k**, Lino, 50 μM), or gabapentin (**l**, Gbp, 300 μM) relative to saline (dashed grey). **m**, Percent change in optogenetic activation by any stimulus train relative to saline as mean and SEM.

While channelrhodopsin-mediated activation of neurons has been used extensively to probe neural circuits, studies examining they contributions of physiologically relevant ion channels to activation have been limited to cell lines overexpressing specific ion channels and not primary neurons with the native cohort of channels(Zhang et al., 2016). Sensitivity of the ChR2 response to different ion channel blockers and activators would support the biological feasibility of leveraging these optically-evoked calcium responses to evaluate mechanisms of neuronal activation as well as to identify blocking drugs (Figure 6g-m). The voltage-gated sodium channel inhibitor tetrodotoxin (TTX, 300 nM) reduced calcium responses to optogenetic stimulation, indicating recruitment of TTX-sensitive sodium channels by optogenetic stimulation (TTX: 57.3±3.9% decrease, n=8 wells/drug)(Blair and Bean, 2002). Voltage-gated calcium channel blockers ω-conotoxin MVIIA and nifedipine both reduced optogenetic activation (ωCtx: 81.0±2.1% decrease; Nif: 31.6±5.9% decrease, n=8 wells/drug), consistent with contributions of N- and L-type calcium channels to sensory neuron activation(Andrade et al., 2010; Barzan et al., 2016; Blair and Bean, 2002; Wang et al., 2000). We then asked whether the method would extend to channels that inhibit neuronal activation. Indeed, we found that the K_V_7 channel activator retigabine and the K_V_7 channel inhibitor linopirdine exerted opposing effects on optogenetic activation, with linopirdine facilitating responses to brief optical trains and retigabine suppressing responses to strong stimuli (Ret: 20.0±7.5% decrease; Lino: 28.6±16.0% increase, n=8 wells/drug)(Brown and Passmore, 2009; Wainger et al., 2014b). These data implicate a wide range of physiologically relevant ion channels in mediating calcium responses to these optogenetic stimuli and suggest the potential power of this approach for identification of novel analgesics. To assess this possibility, we investigated the effect of the well-established chronic pain treatment gabapentin, which acts predominantly by binding α2δ-1 voltage-dependent calcium channel subunits, on optogenetic activation threshold(Sills, 2006). We observed robust inhibition of optogenetic activation (33.8±5.6% decrease, n=8 wells/drug) comparable in amplitude to nifedipine and retigabine, indicating the potential for identification and validation of novel analgesic compounds using this approach.

## Discussion

We developed an experimental platform, termed APPOINT, that combines high-content imaging, robotic liquid handling, and an unbiased analysis pipeline to interrogate the physiology of individual neurons and leverage their physiological profiles to resolve neuronal subtypes. We focused on applications related to nociceptors, on account of the acute need for novel mechanistic insight into pain pathophysiology and non-opioid treatment development(Dahlhamer et al., 2018; Okie, 2010). We identified new components of sensory neuroimmune signaling, including the direct activation of nociceptors by inflammatory mediators as well as human serum, the latter providing a potential novel pain mechanism. Using APPOINT, we developed an approach for high-throughput quantification of optical activation threshold, expanding all-optical physiology options at large scale, and quantify contributions of individual ion channels to calcium flux in neurons.

Contributions of potential pain targets to pain perception depend on molecular context and cell type, and thus traditional target-based screens may be of limited value for analgesic development(Albisetti et al., 2019; Hill et al., 2018; Huang et al., 2018; Rush et al., 2006; Seal et al., 2009). Instead, phenotypic screens, which focus on biologically-relevant phenotypes rather than specific proteins isolated from their biological context, have received increasing enthusiasm from both academia(Wagner and Schreiber, 2016) and industry(Moffat et al., 2017). However, the high-throughput tools required to perform physiology-based phenotypic screens remain limited and flawed(Bradley and Strock, 2019; Sidders et al., 2018). Few physiology-based phenotypic screens of sensory neurons utilizing calcium imaging have been executed, although two used kinetic plate readers for analysis of calcium flux in primary mouse(Brenneis et al., 2014) and iPSC-derived sensory neurons(Stacey et al., 2018). Because the imaging technique could not deliver raw images for subsequent segmentation of individual cells, the utility of this approach for targeted drug discovery remains compromised, due to noise from extracellular regions and the inability to resolve effects of neuronal subtype(Hill et al., 2018). APPOINT resolves these shortcomings.

In contrast to high-throughput physiology approaches, traditional rig-based physiological techniques provide detailed phenotypes of individual cells(Wainger et al., 2015) but are restricted by the small numbers of neurons and compounds that can be studied. For example, in a recent study of low-threshold mechanosensory neurons, 22 mice were necessary to image a total of 27,216 neurons(Giacobassi et al., 2020). Using APPOINT, this would have been achievable with as few as four mice. Beside throughput, rig-based imaging approaches generally harbor bias from the manual choice of imaging field, post-hoc selection of neurons for analysis, and identification of responding cells using only an empirically selected amplitude threshold. Using APPOINT, we have automated each of these components, and taken advantage of additional features of the individual traces besides response amplitude.

One challenge in developing these unbiased approaches was the lack of an unbiased “ground truth” distinguishing true positive and true negative responses. This eliminated the possibility of calculating any accuracy metric using a single model. Thus, we decided an acceptable alternative would be to compare results from two distinct methodologies, unsupervised hierarchical clustering and supervised random forest models. Surprisingly, the two techniques classified >85% of all responses in the same way. Rigorous, manual inspection of response profiles indicated a high degree of accuracy for both positive and negative determinations, with even large amplitude stimulus artifacts correctly being classified as negative. Furthermore, the generalizability of our random forest models across batches indicated that these models reliably incorporated genuine features of calcium responses to stimuli, rather than over-fitting to batch-specific features. We present APPOINT as an integrated method for both *in vitro* calcium imaging and data analysis, however, implementation of the unbiased analysis pipeline alone, which is compatible with traditional rig-based imaging platforms, has advantages for both increasing throughput and reducing bias compared with typical methods.

Some sensory modalities, such as heat(Caterina et al., 1997) and mechanoreception(Coste et al., 2012), have clearly identified stimuli and receptors, whereas the receptors mediating spontaneous pain are unknown. Despite robust investigation of nociceptor sensitization by inflammatory mediators, how nociceptors spontaneously activate during acute and chronic pain remains largely unclear. At the same time, direct activation of nociceptors offers a clear explanation for an increasingly large number of neuroimmune processes(Chiu et al., 2013), and several reports documented metabotropic activation of nociceptors causing pain *in vitro* and *in vivo*(Eskander et al., 2015; Liu et al., 2010; Mørk et al., 2003; Oh et al., 2001).

Combining APPOINT with unbiased *in silico* identification and ranking of candidate metabotropic receptors revealed metabotropic receptor agonists that elicited calcium flux selectively in physiologically-defined subtypes of sensory neurons: for example, neuropeptide Y activated cells that exhibited functional expression of TrpV1 but neither P2×3 nor TrpM8. In contrast, other ligands such as treprostinil and LPA activated more broadly across nociceptor subtypes. These examples further support the importance of subtype in interpreting pharmacological studies of nociceptors(Hill et al., 2018).

Of the metabotropic receptors we identified, most have previously been linked to pain phenotypes, although direct effects on nociceptor physiology may now warrant additional investigation. These established targets include receptors for cytokines, neurotrophic factors, neuropeptides, and inflammatory lipids(Denk et al., 2017; Inoue et al., 2004). Additionally, compounds targeting 11 of the 18 identified metabotropic receptors are either in clinical trials or have already been approved to be used clinically for other indications. Thus, should these targets prove potent in additional *in vivo* validation studies, repurposing can offer a fast and inexpensive path to clinical trials for pain(Avorn, 2015).

Abnormalities of vascular beds contribute to a large number of painful conditions, ranging from common bruises that follow mechanical injury to retroperitoneal and peritoneal hematomas, subarachnoid hemorrhage, migraine, and cerebral and peripheral vasculitides. Although mechanisms for enhancing pain signaling have been demonstrated in specific diseases(Dodick, 2018), the fundamental initiation of pain signaling–how first order nociceptor neurons become activated–remains unclear. As extravascular leakage of blood components may be a unifying feature of these painful conditions, we considered the possibility that serum may directly activate sensory neurons. Indeed, serum elicited large amplitude calcium responses in a large population of sensory neurons, including both capsaicin-sensitive and αβ-meATP-sensitive putative nociceptors. Potentially confounding changes of osmolarity or pH resulting from serum dilution could not have accounted for such strong activation(Liedtke et al., 2000; Viana et al., 2001; Wemmie et al., 2013). Furthermore, the decrease in serum-mediated activation following either proteinase K treatment or depletion of the top 12 abundant serum proteins supports specific activation by a proteinaceous component present in serum. Thus, we propose that regulation of vascular permeability and blood leakage, which occurs through an array of neuroimmune interactions(Pinho-Ribeiro et al., 2017), may be a direct mechanism of nociceptor activation. Although serum injection into healthy volunteers was reported to be painless, factors released from platelets can elicit robust action potential firing in rodent sensory neuron axons, as well as cause pain in healthy individuals(Ringkamp et al., 1994; Schmelz et al., 1997). Furthermore, alterations in serum composition in the context of pathological pain, as well as sensitization of sensory neurons, may increase the pain induced by serum injection in patients. Elucidating the specific ligands and targets responsible for the observed activation will require further work, and *in vivo* studies will be necessary to demonstrate the implications of this mechanism in acute and chronic pain models. We anticipate that APPOINT will now help us to perform large-scale comparisons of nociceptor activation by serum from healthy volunteers and patients with neuropathic and inflammatory chronic pain conditions, potentially leading to use of serum-dependent neuronal activation as an unbiased, physiological biomarker for pathological pain.

Optogenetics has transformed circuit-level neuroscience and led to understanding of how individual cell types function within broad brain circuits and relevant behaviors(Tye and Deisseroth, 2012). However, whether optogenetic activation of neurons can be used to characterize the endogenous channels that determine neuronal excitability has not been previously tested in neurons. A prior elegant study in an HEK cell line equipped with an heterologously-expressed voltage-gated sodium channel and leak potassium channel showed how optogenetic activation could reveal properties of sodium channel modulation(Zhang et al., 2016). Here, we show that weak optogenetic stimulation of primary sensory neurons can prime neuronal activation while preserving sensitivity to endogenous sodium, calcium, and potassium channel modulators. In the example of Kv7 channels, we show sufficient dynamic range to capture both decrease and increase of these conductances. Although we focus on a calcium readout here, we anticipate more closely approximating traditional rheobase measurements by incorporating recent fluorescent voltage indicators that have substantially improved sensitivities(Abdelfattah et al., 2016). Future studies can now examine nociceptor-dependent processes involved in pathological pain such as peripheral sensitization as well as screens to reduce nociceptor activation.

## Acknowledgements

This research was supported by a NIH New Innovator Award (1DP2-NS106664-01), the New York Stem Cell Foundation, and MGH Departments of Neurology and Anesthesia, Critical Care & Pain Medicine (B.J.W.). B.J.W. is a New York Stem Cell Foundation – Robertson Investigator. We thank Marco Loggia for providing samples, Anne-Louise Oaklander and members of the Albers and Lagier-Tourenne labs for helpful discussions.

## Author contributions

All authors reviewed and edited the manuscript. D.M.D. and B.J.W designed experiments and wrote the paper. D.M.D, B.C., K.Z., P.F., X.L., and Y.S. performed experiments. D.M.D. and B.J.W analyzed and interpreted data.

## Competing Interests

The authors declare no competing interests.

## Materials and Methods

### Primary mouse DRG isolation and plating

All animal protocols were approved by the MGH IACUC. Mice (C57Bl6, 2-8 weeks old, male and female) were euthanized by CO_2_ asphyxiation, followed by decapitation. DRG (C1-L6, left and right) were quickly dissected into ice-cold DMEM/F-12 (Thermo Fisher 11320082) and dissociated using a solution of collagenase A (Sigma-Aldrich 10103578001, 2 mg/mL) and dispase (Thermo Fisher 17105041, 2 mg/mL) diluted in Hank’s Balanced Salt Solution (Thermo Fisher 14185052) for 60-90 minutes at 37°C, followed by mechanical trituration using a flame-polished Pasteur pipette. Dissociated cells were filtered using a 70 μm cell strainer (Thermo Fisher 22363548) and BSA gradient (Sigma-Aldrich A9576, 10% in PBS, centrifuge 12 min, 200 rcf). Cells were cultured in neurobasal media (Thermo Fisher 21103049) supplemented with B27 (Thermo Fisher 17504044), GlutaMax (Thermo Fisher 35050061), Pen/Strep (Thermo Fisher 15070063) and AraC (R&D Systems 4520) in 96-well plates (Ibidi 89626) treated with poly-d-lysine(Sigma-Aldrich A-003-E, 1-2 hr at 37°C, 2 μL/well) followed by laminin (Thermo Fisher 23017015, 1-2hr at 37°C, 2 μL/well) overnight at 37°C. All primary sensory neurons were stained and imaged at 1 day in vitro.

For optogenetic activation experiments, Trpv1-Cre (Jax 017769) male mice were crossed with lsl-ChR2-EYFP (Jax 024109) female mice and first-generation Trpv1/ChR2-EYFP pups (male and female, 2-8 weeks old) were used for preparation of sensory neurons.

### Calcium imaging

Primary mouse and human iPSC-derived sensory neurons were stained with the calcium indicator Fluo4-AM for monitoring intracellular calcium concentration. For both cell types, cells were washed once with physiological saline (in mM: 140 NaCl, 5 KCl, 2 CaCl_2_, 1 MgCl_2_, 10 D-glucose, 10 HEPES and pH 7.3-7.4 with NaOH) followed by incubation with Fluo4-AM (Thermo Fisher F14201, 3 μg/mL in 0.3% DMSO) in culture media in the dark at room temperature for 30 min. After 30 min, dye-containing media was removed and replaced with 100 µL of saline and immediately transferred to the imaging chamber. For optogenetics experiments, loading with the calcium indicator CalBryte-630AM (AAT Bioquest 20720, 3 μg/mL in 0.3% DMSO) followed the same protocol.

All calcium imaging experiments were conducted using an ImageXPress micro confocal high content imaging system (Molecular Devices) with automated robotic liquid handling. Cells were maintained at 37°C and with 5% CO_2_/O_2_ during imaging and were imaged (60 s/well) using widefield mode with 10x magnification at a frequency of 1 Hz in the center of each well. Asynchronous liquid dispensing was used for all stimulus applications, allowing for continuous cell visualization during stimulation. All stimuli were a total volume of 20 μL and delivered into the bath solution at a rate of 5 μL/s. Stimulus concentrations are reported as the initial concentration in the stimulus plate, not the final concentration after mixing with the cell bath solution. Doses for all stimuli are indicated in the main text and all stimuli and treatments were diluted in physiological saline, with DMSO (maximum 0.1%) or ethanol (maximum 0.1%) when necessary. Saline stimulation used the same physiological saline, with 0.1% DMSO or 0.1% ethanol when appropriate. DMSO and ethanol did not elicit any direct activation compared with saline alone.

Optogenetic stimuli consisted of five trains of increasing numbers of stimuli (1,3,5,7, and 10 pulses per train) of 1 ms each, delivered at 50 Hz with 10 s between successive trains. CalBryte-630 intensity was measured at 5 Hz.

### Quantification of neuronal activation

All custom analysis scripts will be made available upon request. Initial identification and quantification of Fluo4 intensity was performed using a custom journal in MetaXPress (Molecular Devices) analysis software. Briefly, Flou4-positive cells in each well were identified using a minimum projection of the timelapse image stack based on size and Fluo4 intensity. Each cell automatically generated an individual ROI, which was transferred back to the original timelapse stack and the mean intensity of each ROI was calculated for every image. Raw data by cell for each well were saved as a csv file for further analysis. Non-neuronal cells remaining from initial dissociation and not eliminated by AraC treatment were reliably excluded during cell segmentation based on their substantially smaller cell body size.

Analysis of intensity and quantification of neuronal activation was performed in R (v3.5.0). Briefly, raw intensity values for each cell were normalized to the baseline intensity of the initial 5 images. Peak response amplitude was calculated as the median of 3 intensity values centered on the peak intensity within 15 s after stimulus onset, with independent baselines of the 5 s preceding stimulus onset for each stimulus. Maximum rate of rise of Fluo4 was calculated as the peak of the derivative of intensity values within 15 s after stimulus onset.

For hierarchical clustering analysis, pooled responses were filtered to center clustering around typical thresholds (peak amplitudes between 0.075 and 0.225 dF/F; maximum rate of rise between 0.5 and 1.5 dF/F/s). Hierarchical clustering used the euclidean distance between observations and Ward D2 agglomeration algorithm. Random forest modeling was accomplished using caret (v6.0) and randomForest (v4.6) R packages using default settings and selecting the simplest model within one standard error of the optimal model.

### Human iPSC differentiation

Induced pluripotent stem cells (iPSCs) of a single line, 446, from a healthy individual(Lee et al., 2009) were obtained under MGH IRB approval and differentiated into sensory neurons according to an established protocol(Schwartzentruber et al., 2018). Frozen stocks of a single passage were thawed and expanded in Matrigel (Corning 354277) coated 6-well plates (Corning 353046) for 3 days using mTESR1 media (Stem Cell 85850) before initiating differentiation. On day 1 to day 3 of differentiation, cells received hESC media [DMEM/F12 media with Knock-out Serum Replacement (Thermo Fisher 10828-028), non-essential amino acids (Corning 25-025-CI), GlutaMax, and β-mercaptoethanol (Thermo Fisher 31350010)] supplemented with LDN-193189 (Stemgent 04-0074-02) and SB431542 (DNSK International DNSK-KI-12). Beginning on day 4, hESC was supplemented with the combination of SU5402 (DNSK International DNSK-KI-11), CHIR 99021(Tocris 4423), and DAPT (DNSK International DNSK-EI-01), in addition to LDN and SB. For days 5-10, NB media [Neurobasal with N2 (Thermo Fisher 17502048), B27, GlutaMax, and β-mercaptoethanol] was increasingly added to to hESC media every 2 days, such that media for days 5 and 6 constituted 75% hESC/25% NB, for days 7 and 8 constituted 50% hESC/50% NB, and for days 9 and 10 25% hESC/75% NB. Supplementation with SU, CHIR, and DAPT was maintained during this transition, but LDN and SB were removed after day 7. On day 11, immature neurons were dissociated using accutase (Worthington LK003178) and replated at a density of 3*10^6^ cells/well in Matrigel-coated 6-well plates containing NB media supplemented with BDNF (Thermo Fisher PHC7074), GDNF (Thermo Fisher PHC7044), NGF (R&D 256-GF), NT-3 (Thermo Fisher PHC7036), and ascorbic acid (Sigma-Aldrich A4403). Media was changed twice weekly thereafter until 32 or 63 days *in vitro* for staining and physiology, respectively.

### Immunohistochemistry

iPSC-derived sensory neurons were dissociated at 32 days *in vitro* using papain for 25-35 min and replated into 96-well plates (Ibidi 89626). They were maintained in NB supplemented with BDNF, GDNF, NGF, NT3 and ascorbic acid for 3 days, followed by fixation with 4% paraformaldehyde (Thermo Fisher 50-980-495). Cells were blocked for 4 hrs at 4°C, incubated with primary antibodies for Isl1/2 (mouse, DSHB 39.4D5, 1:500) and Peripherin (rabbit, Millipore AB1530, 1:200) overnight at 4°C, followed with donkey anti-mouse (Thermo Fisher a21202) and donkey anti-rabbit (Thermo Fisher A10042) secondary antibodies at 23°C for 4 hours. Cells were imaged using ImageXPress micro confocal high content imaging system (Molecular Devices).

### ASO design

ChR1, ChR2, ChR3 and scrambled ASOs (Integrated DNA Technologies) were constructed using a gapmer design, phosophorothioate bonds (indicated by * in sequence), and 2’-O-methyl modified RNA bases (indicated by preceding “m”) flanking unmodified DNA bases (indicated by underline). ASO sequences are listed in Table 1 and all ASOs were applied at 10 μM for 7 days to sensory neurons on day 0 of dissociation. Culture media was supplemented with NGF to promote neuronal health during extended culture.

**Table 1.** ASO sequences.

| ASO | Sequence |
| --- | --- |
| ChR1 | mU*mU*mG*mU*mA*<u>C*T*C*C*A*G*C*T*T*G</u>*mU*mG*mC*mC*mC |
| ChR2 | mC*mC*mA*mU*mG*<u>G*G*T*C*T*G*C*T*T*G</u>*mU*mG*mU*mC*mU |
| ChR3 | mC*mU*mC*mA*mG*<u>G*T*A*G*T*G*G*T*T*G</u>*mU*mC*mG*mG*mG |
| Scrambled | mA*mG*mC*mC*mC*<u>C*A*C*A*T*C*T*T*T*G</u>*mC*mU*mA*mC*mC |

### In silico analysis of transcriptomic datasets

Read count data for GSE10246, GSE55114, GSE63576, and GSE89224 were downloaded from GEO and loaded into R (v3.5.0). The data were filtered to include only genes present in all four datasets and fold change and adjusted p-values from each dataset were pooled. Mean expression counts from GSE63576 were used to reflect relative expression strength in mouse DRG. Genes were then filtered to identify relevant genes with expression regulated by nerve injury (adjusted p-value spared nerve injury v sham<0.001), or preferential expression in DRG relative to other tissues (adjusted p-value DRG v Other<0.005 and expression > 10), or preferential expression in nociceptors relative to proprioceptors (adjusted p-value Nociceptors v Proprioceptors<0.002 and expression >3). Metabotropic receptors were identified using all genes associated with gene ontology terms (0030594, 0005030, 0038187, 0001653, 0004888). Each remaining gene was then ranked using the weighted average of fold-change enrichment and raw expression (weights: SNI enrichment=0.3, Nociceptor enrichment=0.3, DRG enrichment=0.2, raw expression=0.2).

### Serum-dependent activation

Blood samples were collected from healthy individuals (5 male and 4 female, 25 to 85 years old, mean age=47 years) under MGH IRB approval. Serum was prepared and frozen in aliquots at −80°C. For activation, serum was thawed and split into four portions for: 1) unmodified serum stimulation; 2) heat inactivation at 56°C for 1 hr; 2) proteinase K digestion (New England Biolabs P8107S) at 37°C for 18 hrs; 3) depletion of top 12 abundant serum proteins (Thermo Fisher 85165) according to manufacturer instructions. The 12 most abundant serum proteins include: albumin, IgG, α1-acid glycoprotein, α1-antitrypsin, α2-macroglobulin, ApoAI, ApoAII, fibrinogen, haptoglobin, IgA, IgM, transferrin.

### Statistical Analysis

Mean and standard error of the mean are reported throughout, unless specified. Details of specific statistical analyses are included in the main text. For differences between cumulative distributions, we used the two-sample Kolmogorov-Smirnov test of the hypothesis that both individual distributions are drawn from the same underlying distribution. Significant changes in activation are calculated as percent change from baseline or vehicle condition by batch and significance assessed as one-sample t-test of the hypothesis that the net change across batches is zero. Batch effects are assessed using univariate ANOVA, followed by Tukey HSD correction for multiple comparisons when appropriate.

### Data Availability

The datasets generated and/or analysed during the current study are available from the corresponding author on reasonable request

### Code Availability

All data analysis scripts will be made freely available from the corresponding author on request.

## Notes

### Competing Interest Statement

The authors have declared no competing interest.

