## Supplemental figures for "A high-content platform for physiological profiling and unbiased classification of individual neurons"

**Supplementary Figures 1-6**

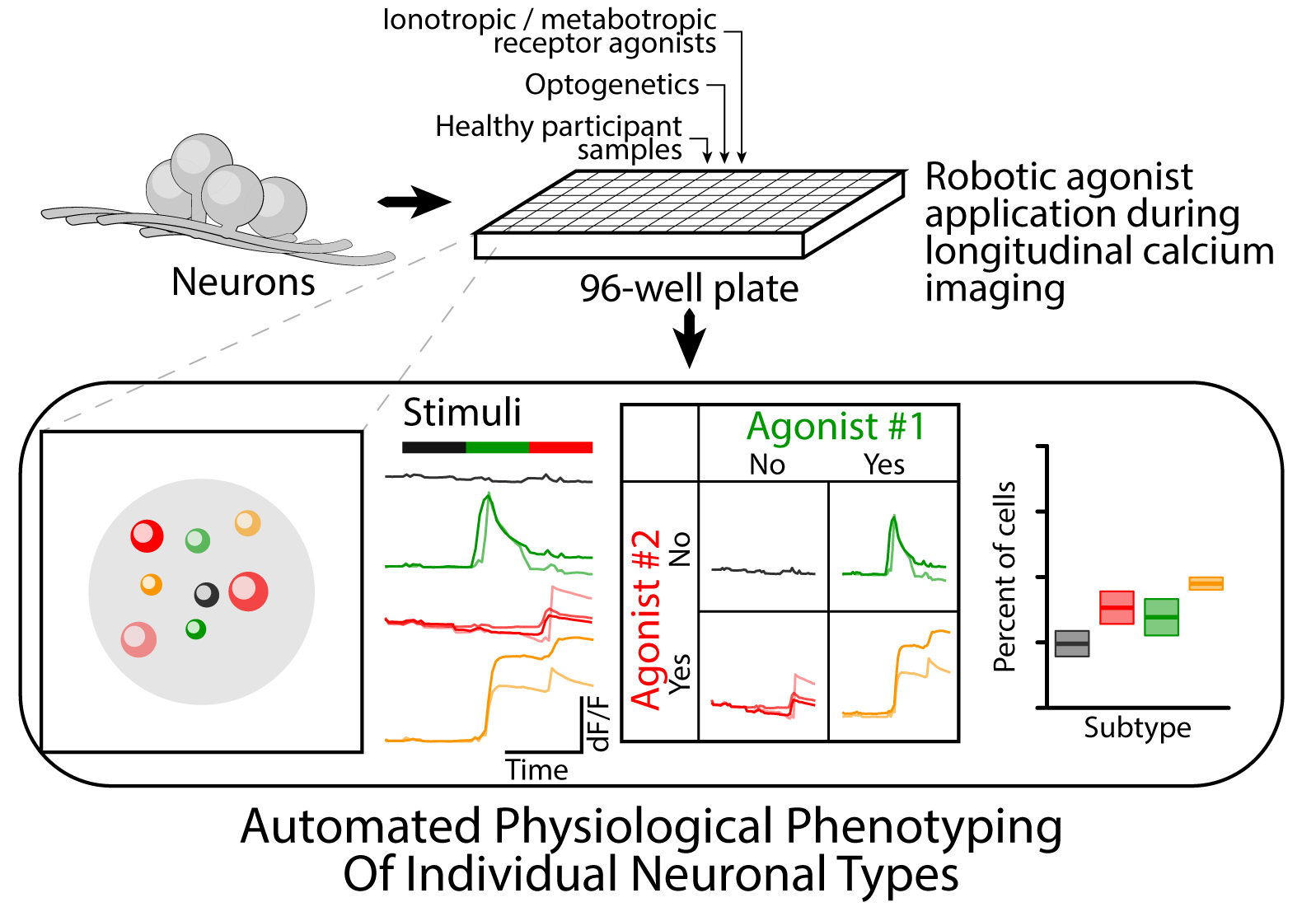
**Supplementary Figure 1. Overview of APPOINT**. Neurons are plated into 96-well format and stimulated by automated application of sequential chemical or optical stimuli during longitudinal calcium imaging. Individual neurons are automatically segmented and response profiles for each series of stimuli are extracted. Based on the unbiased classification of responses to each agonist, individual neurons are sorted into distinct functional subtypes and the frequency of each subtype is quantified.

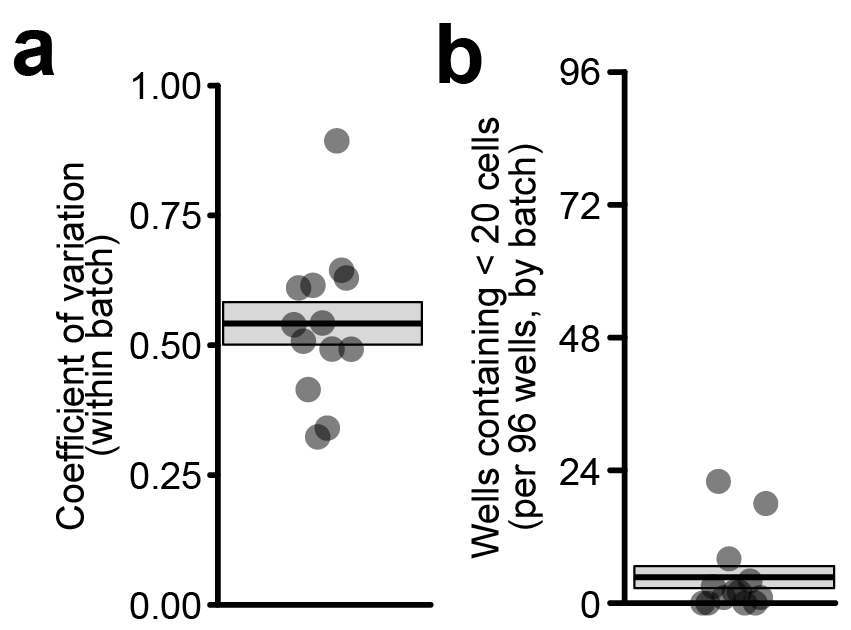
**Supplementary Figure 2. Neurons are evenly distributed across individual plates with few low-density wells.**

**a,** Coefficient of variation within each plate (points), along with mean and SEM (box, 0.542 ± 0.041).

**b**, Wells per 96-well batch with fewer than 20 cells per well (points), along with mean and SEM (box, 4.7 ± 2.0 wells per 96-well batch).

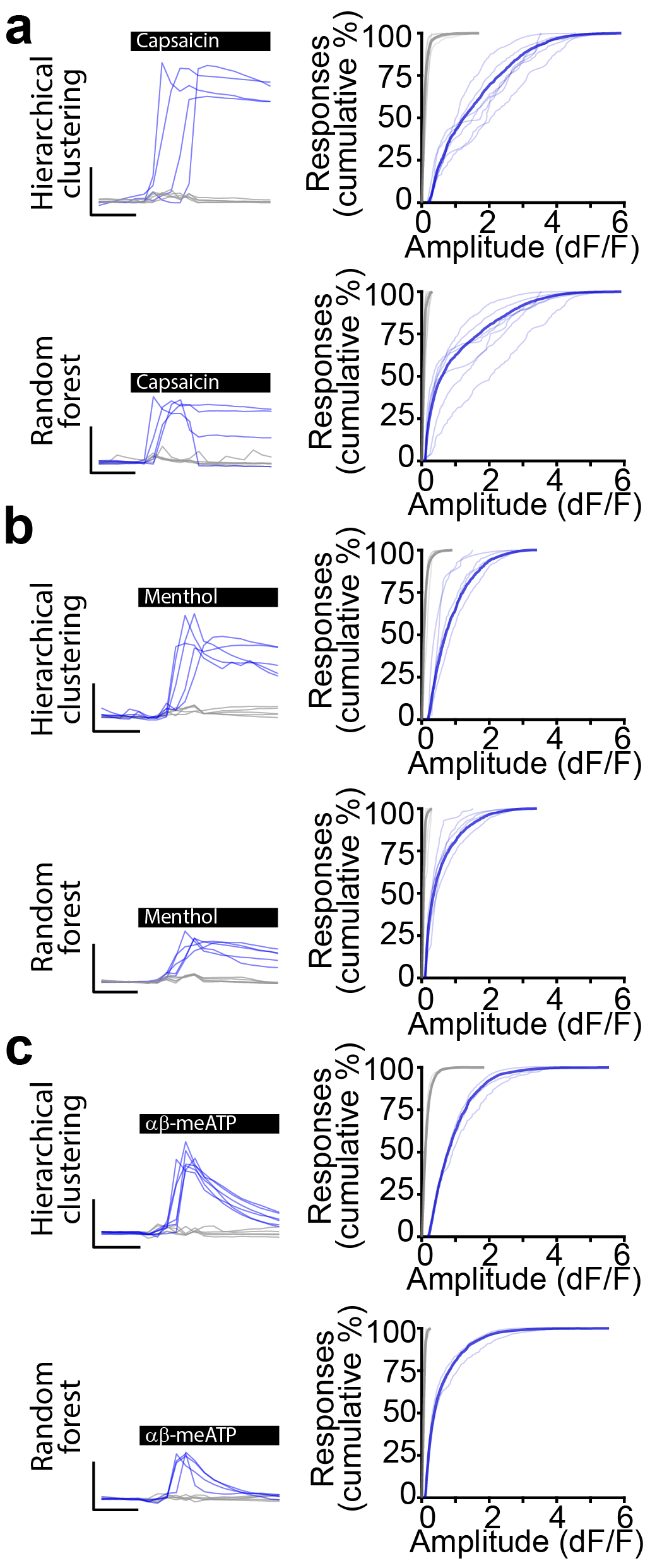
**Supplementary Figure 3. Unbiased classification of responses to Trp and puringergic receptor agonists.** Classification of calcium responses following application of **a,** capsaicin**, b,** menthol**,** and **c,** meATP by hierarchical clustering (top) and random forest machine learning (bottom) approaches. Each panel shows example traces of calcium responses classified as positive (blue) and negative (grey) (left, scale bars 5 s and 0.5 dF/F) along with cumulative distributions of positive (blue) and negative (grey) response amplitudes (right).

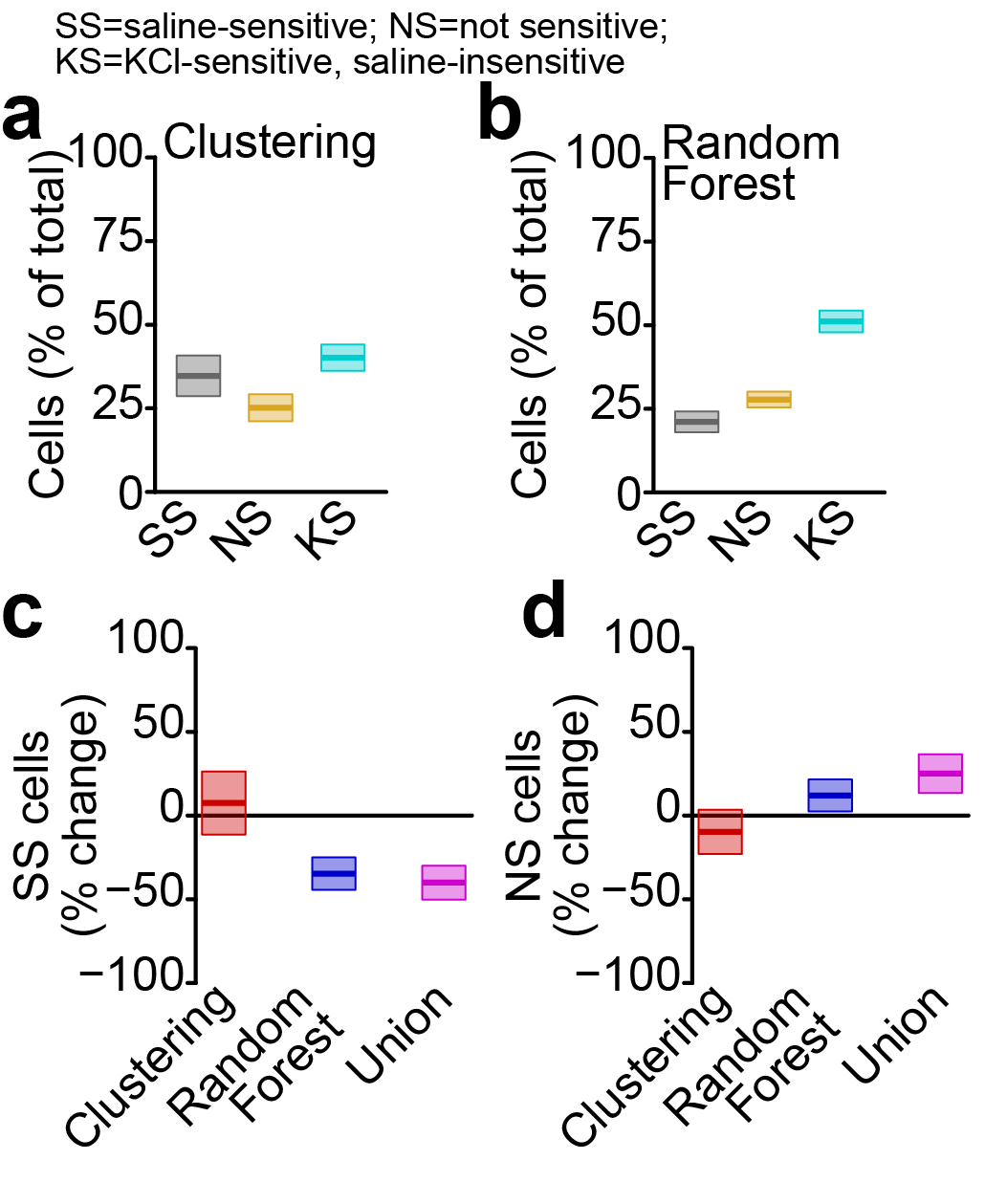
**Supplementary Figure 4. Comparison of individual, unbiased approaches for quantification of cell types. a-b,** Percentage of cells classified as SS (grey), NS (orange), and KS (cyan) using hierarchical clustering (**a**) or random forest classifiers (**b**). **c-d,** Percent change from threshold-based analysis in percentage of SS cells (**c**) or NS cells (**d**) using hierarchical clustering (red) or random forest (blue) classifiers alone or the union of both approaches (magenta).

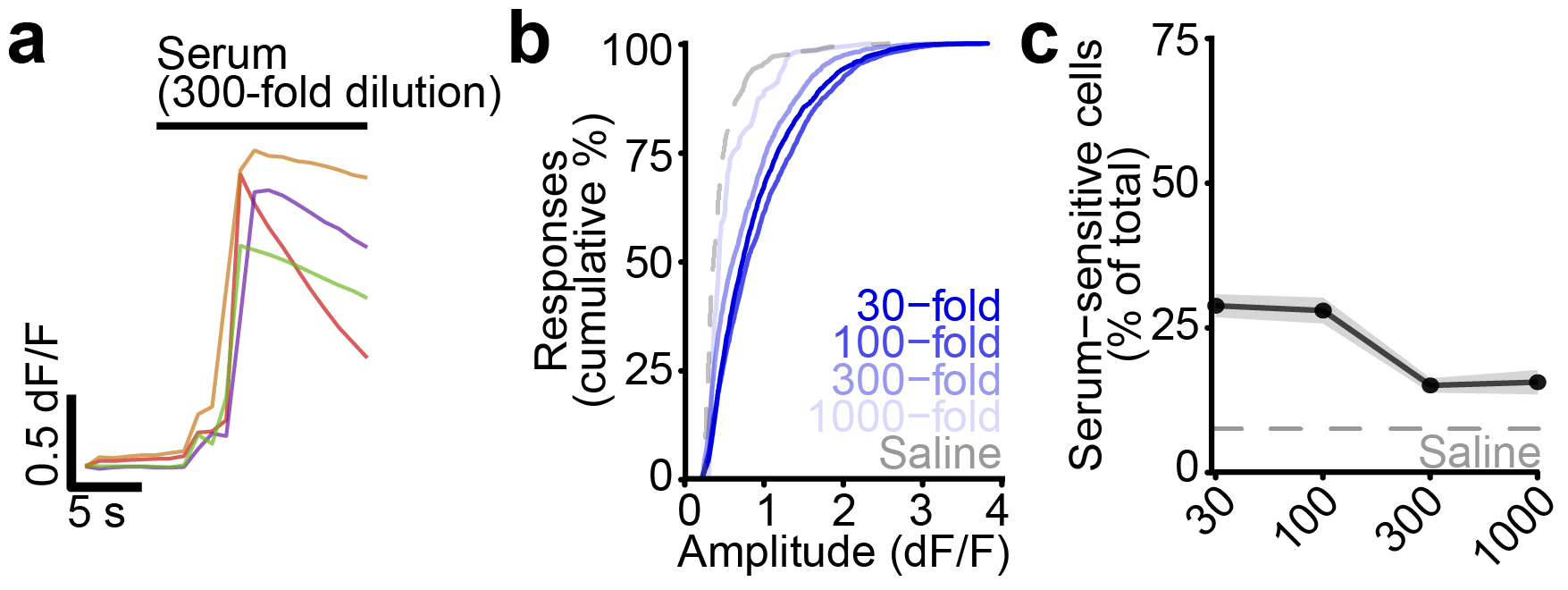
 **Supplementary Figure 5. Activation of primary mouse sensory neurons by human serum samples.**

**a,** Example traces of positive responses to serum (300-fold dilution, colors indicate individual cells, scale bars 5 s and 0.5 dF/F).

**b**, Cumulative distributions of positive response amplitudes following stimulation by serum dilutions (as fold dilution, orange), along with meATP (dashed green line) and saline (dashed grey line). Analysis includes cells sensitive to initial saline stimulus.

**c**, Dose-response curve of sensory neuron activation by serum as mean (points/lines) and SEM (shaded area), along with saline-dependent activation (dashed line). Analysis includes cells sensitive to initial saline stimulus.

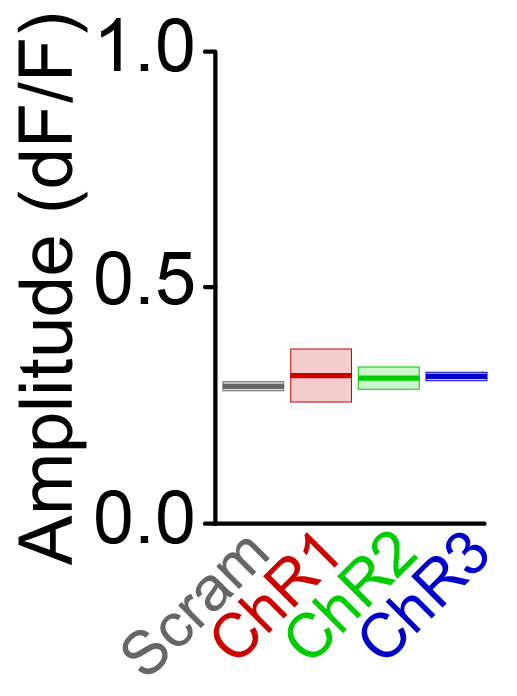
**Supplementary Figure 6. Effect of ChR-targeting ASOs on amplitude of optogenetic responses.** Average response amplitude of positive responses following optical stimulation after seven-day treatment by ChR2-targeting ASOs (red, green, blue) or scrambled control ASO (grey) as mean and SEM.

**Supplementary Tables 1-2**

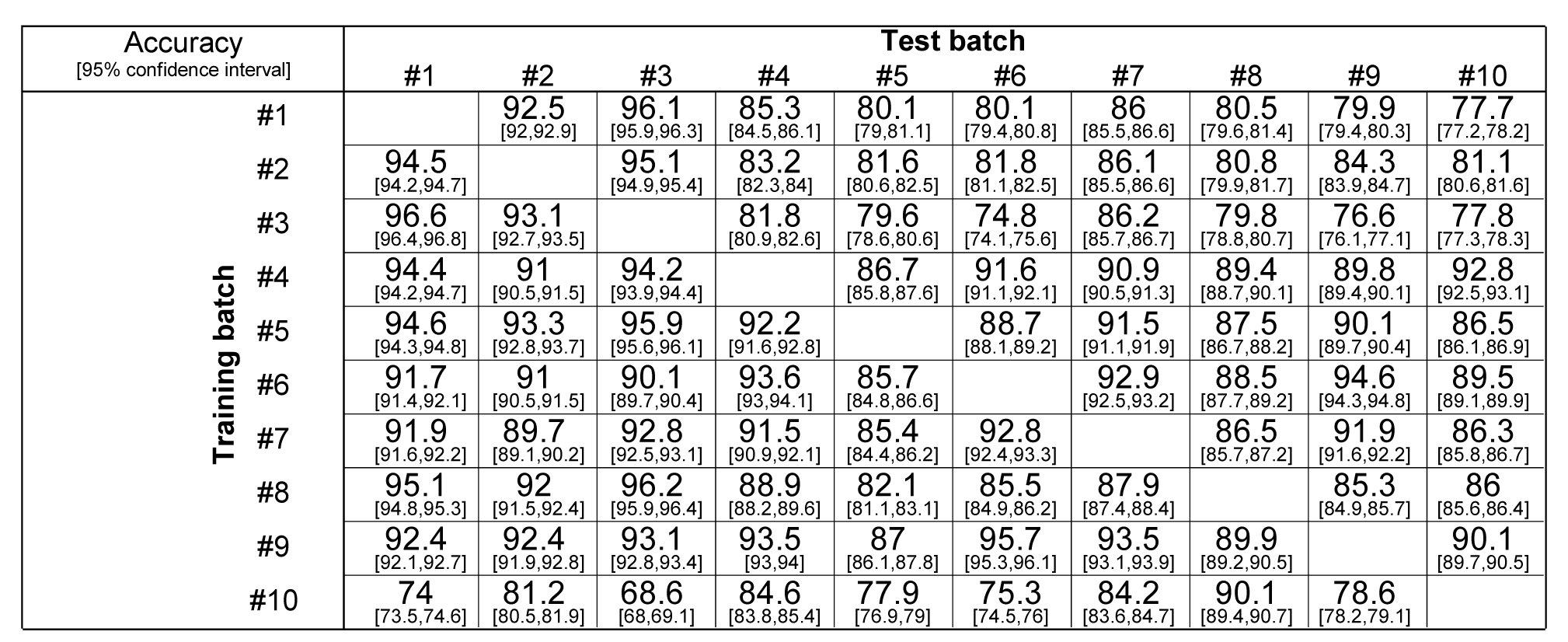
**Supplementary Table 1. Robust classification of responses across neuronal batches.** Accuracy and 95% confidence intervals for pairwise classification of individual test batches using random forest models trained on independent single batches. All accuracy values are statistically significant with p < 5*10^-100^.

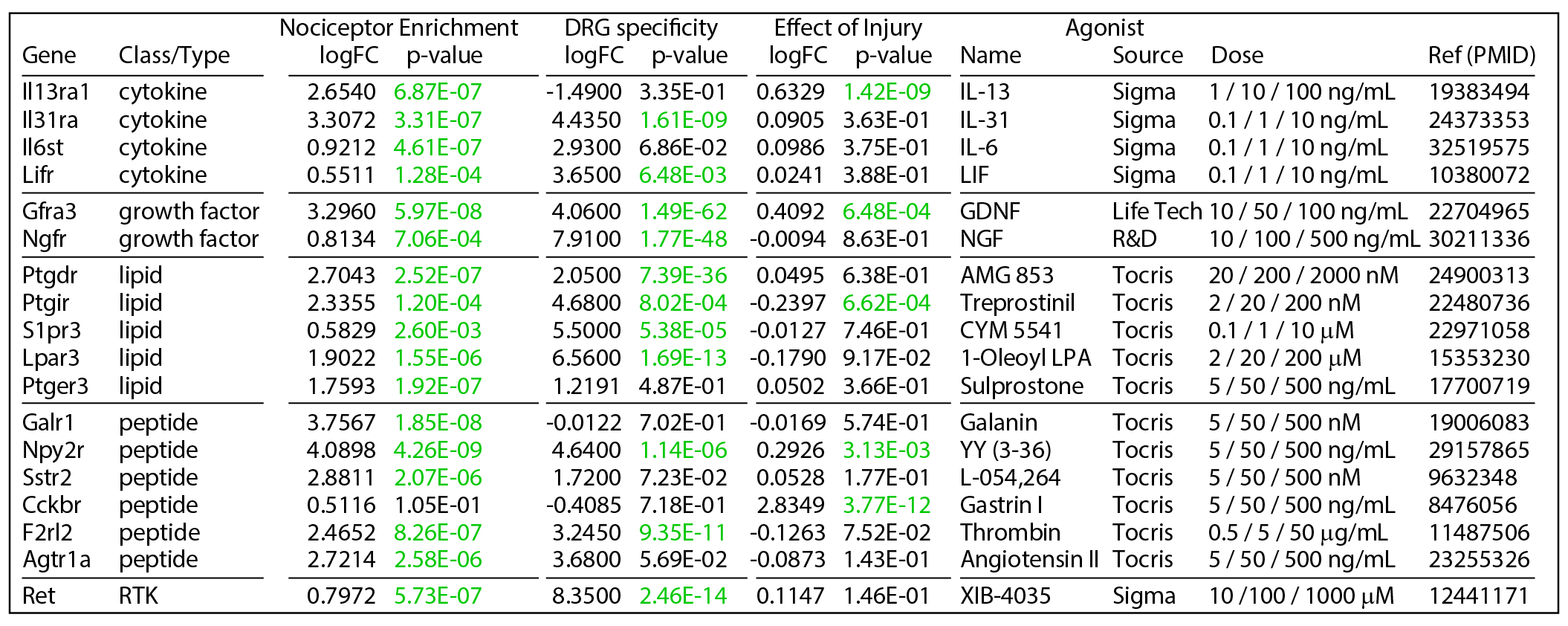
**Supplementary Table 2. Pool of metabotropic receptor agonists from *in silico* analysis of transcriptomic datasets.**
